# Investigating the conformational landscape of AlphaFold2-predicted protein kinase structures

**DOI:** 10.1101/2022.12.02.518928

**Authors:** Carmen Al-Masri, Francesco Trozzi, Marcel Patek, Anna Cichońska, Balaguru Ravikumar, Rayees Rahman

## Abstract

Protein kinases are a family of signalling proteins, crucial for maintaining cellular homeostasis. When dysregulated, kinases drive the pathogenesis of several diseases, and are thus one of the largest target categories for drug discovery. Kinase activity is tightly controlled by switching through several active and inactive conformations in their catalytic domain. Kinase inhibitors have been designed to engage kinases in specific conformational states, where each conformation presents a unique physico-chemical environment for therapeutic intervention. Thus, modeling kinases across conformations can enable the design of novel and optimally selective kinase drugs. Due to the recent success of AlphaFold2 in accurately predicting the 3D structure of proteins based on sequence, we investigated the conformational landscape of protein kinases as modeled by AlphaFold2. We observed that AlphaFold2 is able to model several kinase conformations across the kinome, however, certain conformations are only observed in specific kinase families. Furthermore, we show that the per residue predicted local distance difference test can capture information describing conformational dynamics of kinases. Finally, we evaluated the docking performance of AlphaFold2 kinase structures for enriching known ligands. Taken together, we see an opportunity to leverage AlphaFold2 models for structure-based drug discovery against kinases across several pharmacologically relevant conformational states.

## 1 Introduction

Protein kinases are a family of more than 500 proteins that catalyze the transfer of a phosphate group to their substrates, switching specific cellular pathways on or off. As a result of mutations, differential expression or other forms of dysregulation, kinases are known to cause diseases such as cancer and autoimmunity (1; 2). Consequently, they are one of the largest drug target families in the druggable genome (3). Importantly, kinases are structurally dynamic proteins that can adopt several conserved active and inactive conformational states. These specific conformations regulate important aspects of cellular physiology and are key driving factors for protein-protein (4) and protein-ligand interactions (5).

In the active conformation of protein kinases, the conserved DFG motif and *α*C helix have an “in” conformation (CIDI). This means that the conserved phenylalanine of the DFG motif (DFG-Phe) points out of the active site while the aspartate (DFG-Asp) faces the ATP-binding site. Additionally, the conserved glutamic acid in the center of *α*C helix (C-Glu) forms a salt bridge with the conserved *β*3-lysine. Conversely, inactive conformations comprise of either the DFG or *α*C helix adopting the “out” conformation (CIDO, CODI, CODO) (6). That is, the directionality of the DFG-Asp and DFG-Phe are flipped, or the glutamic acid in the *α*C helix breaks contact with *β*3-lysine. Additionally, an inactive conformation can arise when the DFG-Phe adopts an intermediate conformation (*ω*CD) (6).

Inhibitors binding to these distinct conformational states are also well-characterized. For example, type I inhibitors are ATP competitive molecules that bind to the active CIDI conformation; whereas type I1/2 and type II inhibitors engage the inactive CODI or CIDO conformations, respectively (7). These distinct inhibitor types occupy diverse regions of the chemical space and confer pharmacokinetic/dynamic advantages (7). Therefore, approaches that accurately model the different conformational states of kinases can enable the rational design of novel, conformation-specific inhibitors.

Of the 3,700 crystal structures of human protein kinases in the Protein Data Bank (PDB) (8), roughly half of the human kinome is covered (6). Additionally, over 60% of crystal structures catalogue kinases in just their active conformation (9). Due to the striking success of AlphaFold2 (AF2) in the prediction of protein structure from sequence (10), and AF2’s accuracy in modeling membrane transporter conformational diversity (11), we hypothesized that AF2 may be able to model the conformational landscape of protein kinases. In this work, we analyzed all of the modeled protein kinase structures from the AlphaFold Protein Structure Database (AlphaFoldDB) (12). We then evaluated their utility in structure-based drug discovery by calculating the enrichment of known actives to selected kinases using molecular docking. The enrichment observed in the AF2 structures was then compared to the best matched native crystal structure from the PDB.

## 2 Methods

### 2.1 Acquisition and prediction of conformation of PDB and AF2 protein kinase structures

All protein kinase structures present in both the PDB (13) and AlphaFoldDB (https://alphafold.ebi.ac.uk/) were recovered. Using the SCOPe classification (14), only chains containing the “Protein kinases, catalytic subunit” family were retained from the PDB, and the catalytic domain was subsequently extracted from these chains. This filtering process resulted in 5752 unique PDB IDs from which 8460 chains were selected.

From AlphaFoldDB, only the structures containing a “Protein Kinase” domain annotated by UniProtKB (15) were retained. The catalytic domains of these structures were extracted. After this initial filtering step, 4348 structures remained.

Kinformation (6; 16) was used to annotate the different conformations of kinase domains. Kinformation is a random forest model which uses descriptors related to the *α*C helix and DFG motif to predict the conformation of kinases. Kinformation was set to discard sequences with a length less than 225 residues, and less than 40% sequence identity to all canonical human kinases. Structures that passed these filtering criteria were subsequently aligned with the Modi-Dunbrack alignment (17) using MUSCLE (18), and the alignment was further refined through structural alignment using pyMOL (19), with 1ATP as reference.

### 2.2 Binding pocket comparison of AF2-modeled kinase structures

The structures predicted by AF2 were clustered by binding pocket similarity. The binding pocket of each kinase was characterized by a set of 85 residues, as defined by KLIFS (20). These residues characterize the ligand-receptor interactions of type I, I1/2, II, III, and most type IV inhibitors. The 85 residues sequence for each kinase was aligned to their corresponding AF2 structure using Bio.pairwise2(21) (global sequence alignment with a match score of 1, a mismatch penalty of −1, a gap penalty of −4, and no gap extension penalty).

After obtaining the binding pocket residues, we utilized the KiSSim package (22) to extract spacial properties of each pocket residue (default settings). The spacial properties consisted of the distances of each residue to the center of the pocket, to the hinge region, to the front pocket, and to the DFG region. The structures were clustered by using t-SNE (23) (5000 iterations, perplexity of 30, 3 components).

### 2.3 Molecular docking and crystal structure selection workflow

Docking enrichment calculations were carried out for three kinases: ABL1, BTK, and FLT3. For each kinase, a set of representative structures was selected as a result of the following steps: 1. All holo-structures were initially considered; 2. Structures with missing or mutated residues in the binding pocket or had a resolution >2.5 Å were excluded; 3. Each crystal structure was assigned a conformational state following the scheme proposed by Ung et al. (6), and featurized using fpocket(24) along with KLIFS descriptors for the C-helix, DFG, and Glycine rich regions (OpenCADD-KLIFS) (25); 4. These structures were clustered using DBSCAN (26). The representative structure for a given conformation cluster was determined to be the structure that maximized the total number of protein-ligand interactions; 5. Finally, the representative structures were subjected to energy minimization using MOE (27).

These structures were evaluated against benchmark sets of known actives, inactives, and decoys built for each kinase target. A subset of active molecules with known activity <1000 nM (IC50, K*i*, or K*d*) were selected using a diversity picker implemented in RDKit (28) (20% of known actives with < 0.20 average Tanimoto similarity). All known inactives with activity > 9999 nM were selected, and decoys were generated using DeepCoy (29) pre-trained on the DUDE model to achieve a total of 25 negative examples per active. The benchmark set was neutralized using RDKit, and the 3D conformers were generated using openbabel (30).

The docking calculations were performed using Smina (31). Prior to docking, all representative protein structures were superimposed, and all orthosteric ligands were considered to define the consensus binding site. Smina default scoring function and parameters were employed. For each kinase, an early enrichment analysis was performed on structures belonging to the same conformational state of the AF2 structure. The structures with the best enrichment were then selected to be compared to the AF2-generated ones. To calculate the docking enrichment curves, the docked ligands were ranked based on their binding affinity. The ratio of known actives against the total number of compounds was plotted to test the capability of each structure to discriminate actives from inactives and decoys. Lastly, the ability of AF2 structures to reproduce pharamacologically relevant interactions was assessed via cross-docking analysis on a set of 30 kinases. To evaluate crystal structures, each crystal ligand was docked to all crystal structure of the same kinase, except for its native structure. For each AF2 structure, all crystal ligands of the same kinase were docked to it. The reproduction of interactions was assessed by computing protein ligand interaction fingerprints using the protein-ligand interaction profiler package (plip) (32) and compared to native ligand contacts using a Jaccard similarity score.

## 3 Results

### 3.1 The conformational landscape of protein kinase models generated by AF2

To investigate the conformational landscape of protein kinases, we predicted the different conformational states across both human and non-human protein kinase structures present in the RSCB PDB and the AlphaFoldDB databases. We observe that the conformational diversity of AF2-modeled protein kinase structures closely mirrored the proportion of kinase conformations in the PDB (Fig. 1A). Here, the active kinase conformation, CIDI, is highly over-represented in the both AF2-modeled structures and the PDB (69% and 68%, respectively), followed by the CODI conformation (20% and 17%). Interestingly, CIDO structures are under-represented in AF2 compared to the PDB structures. This observation is also reflected in related work by Modi and Dunbrack, where a they observe a large proportion of AF2 models of the human protein kinome are in the DFG-in active conformation (70.8%) while inactive DFG-out structures are underrepresented (33).

**Figure 1:**
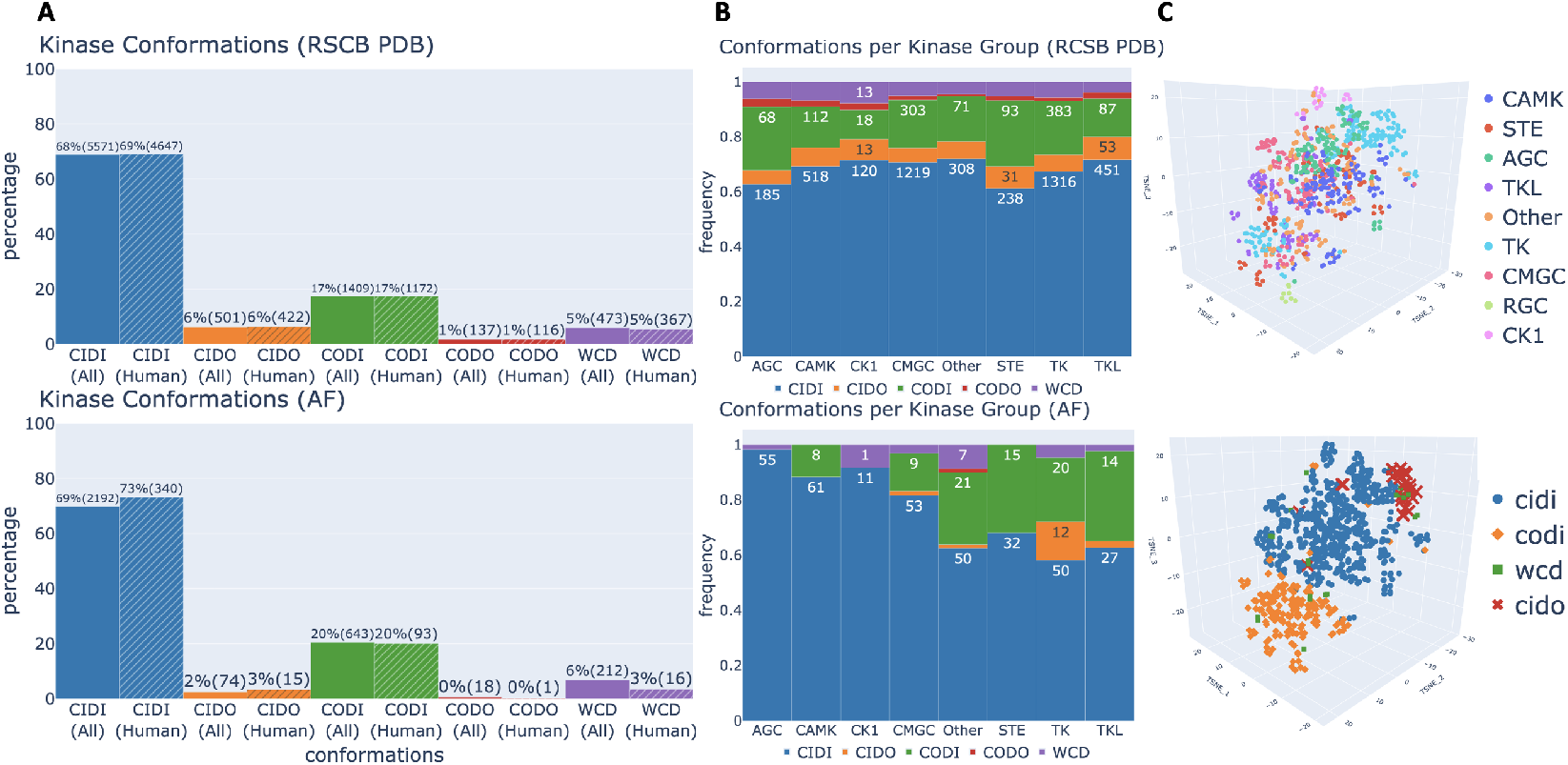
The conformational landscape of protein kinases. (A) The conformational diversity of structures in the RCSB PDB database (top) and AlphaFoldDB (bottom) across all species (non-dashed bar) and Human (Homo Sapiens) species (dashed bar). (B) The proportion of each conformation across kinase families in the human kinome. (C) Clustering of all human and mouse kinase structures using t-SNE, colored by kinase family (top) or conformation (bottom).

The representation of kinase conformation is also dependent on the protein kinase family, where certain kinase families enrich for specific conformational states (Fig. 1B). In the PDB, we observe a relatively consistent proportion of CIDO (type II) kinase structures across all kinase families, as well as other rarer conformations, such as CODO and *ω*CD. However, cross-family conformational diversity is not necessarily conserved in the AF2 models. Specifically, CIDO and CODO conformations are not observed in AGC, CAMK, CK1 and STE families. However, in TK, the relative proportion of CIDO structures is higher in the AF2 models than the PDB structures. We hypothesize that this may be due to a larger diversity of ligands available for TK family kinases that stabilize the type II conformation.

We also investigated the differences in binding pockets across all human and mouse kinases in AF2 models by considering the distances of each of the residues belonging to the pocket to the center, hinge, DFG, and front pocket regions of the kinase (Fig. 1C). After clustering using t-SNE, we found that kinases belonging to the same family form clear sub-clusters (Fig. 1C, top). Many of the kinase families form 2 to 3 sub-clusters, most notably TK, TKL, and CAMK. Only the “Other” family does not cluster together and is spread across the t-SNE projected conformational space. We also find that these observed sub-clusters can be attributed to different conformations of kinases within the same family (CIDI, CODI, and CIDO) (Fig. 1C, bottom), with the structures in the intermediate conformation *ω*CD spread across the t-SNE space.

### 3.2 pLDDT as a measure of conformational plasticity in the protein kinase active site

Each residue of any AF2-modeled structure has an associated estimate of the confidence for its predicted 3D positioning, called predicted local distance difference test (pLDDT). Previous work by Binder et. al has shown that pLDDT scores can correlate to structural properties, such as protein disorder (34). Hegedűs et. al and Guo et al. have also shown a relationship between pLDDT and protein conformational dynamics(35; 36). Since AF2 can model several pharmacologically-relevant kinase conformational states (Fig. 1), we hypothesized if pLDDT correlated with the conformational space sampled by the available crystal structures for each kinase. For example, ABL1 currently has 27 human and mouse structures in the DFG-out conformation, 9 structures in the DFG-in conformation, and 16 structures in an intermediate conformation.

We visualized AF2-modeled structures of four kinases: ABL1, BTK, DDR1, and EGFR. These kinases belong to different kinase families and are representative of multiple conformational states of the binding pocket. We observed variability of pLDDT scores in conformationally flexible regions of the pocket, specifically, in the relative positioning of the residues belonging to the DFG motif (residues 82-84) (Fig. 2A, bottom). We then evaluated the binding pocket pLDDT scores across all AF2 kinase structures and observed that DFG-Phe and C-Glu motifs (residues 20 and 83 in Fig. 2B), have higher uncertainty on average (~ 75%) compared to the rest of the protein.

**Figure 2:**
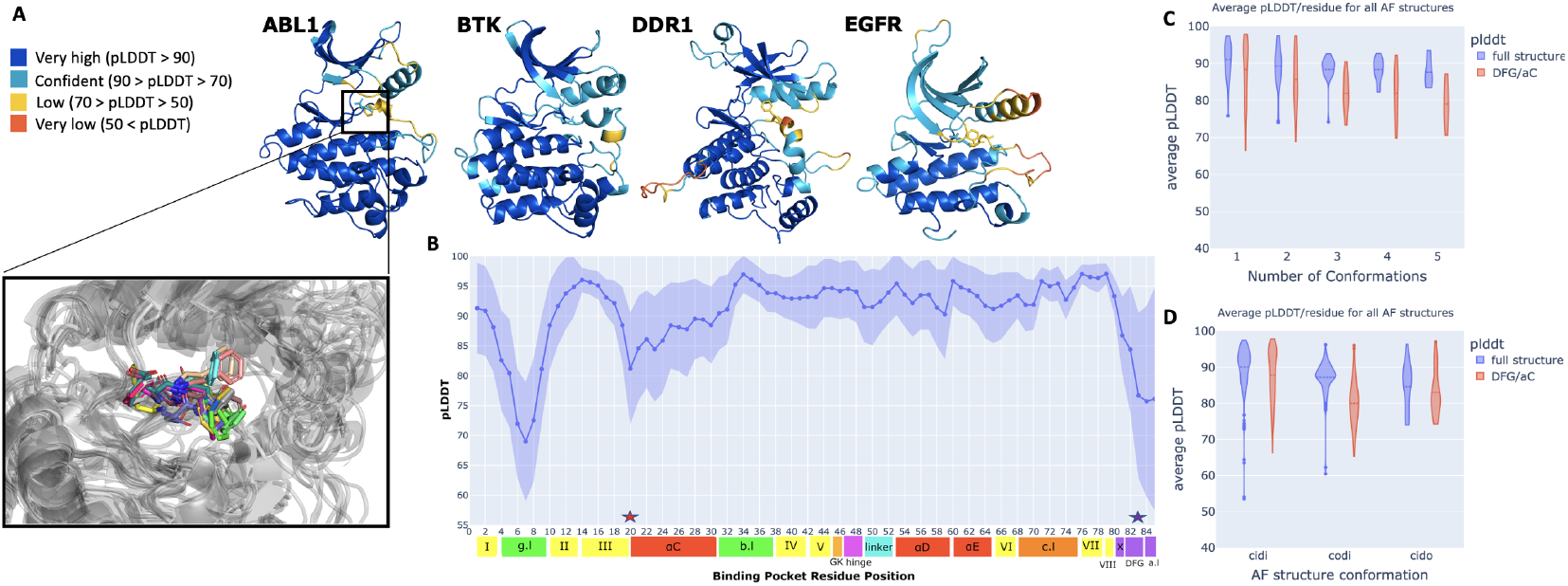
pLDDT as a measure of conformational plasticity. (A) pLDDT for all residues belonging to the protein kinase domain of ABL1, BTK, DDR1 and EGFR. The DFG motif is shown in licorice representation. The variation of the DFG-Phe residue across all human ABL1 structures is also shown (A, bottom). (B) The average pLDDT of the 85 binding pocket residues across all AF2-predicted structures. The standard deviation in the pLDDT is shown as the shaded region. (C) violin plot of the distribution of the average pLDDT across kinases, grouped by the number of conformations available for each kinase. (D) violin plot of the distribution of the average pLDDT across kinases, grouped by the conformation of each structure. In both (C) and (D), the average pLDDT was computed across all the residues in the structure (blue) or across the residues part of the DFG motif and *α*-C helix (red).

We then evaluated if the variability of pLDDT scores in the binding pocket was either related to the number of unique conformations observed in the PDB for each kinase, or was a function of certain underrepresented conformational states, such as CIDO. Figure 2C shows a clear relationship between the average pLDDT of either the whole kinase domain and DFG/*α*-C helix motifs to the number of unique conformations observed in the PDB of a given kinase structure (Fig. 2C). Likewise, underrepresented kinase conformations on average have lower pLDDT in both their whole kinase domain structure and just their DFG/*α*-C helix motifs (Fig. 2D).

### 3.3 Comparison of ligand enrichment in selected, conformationally diverse AF2 kinase models against matched crystal structures

AF2 opens the door to structure-based drug discovery for targets without known crystal structures. To evaluate if these predicted structures can be leveraged for virtual screening, we compared the docking enrichment of the predicted AF2 structures, to the highest scoring minimized crystal structure in terms of docking enrichment for three conformationally diverse kinases (ABL1, BTK and FLT3). We observe that the best enrichment is found for ABL1, predicted in the CIDI state (Fig. 3A). Importantly, the AF2 model performs slightly better than best the holo structure in the early enrichment of known actives (pAUC: 252.54 vs 249.44). This is surprising considering that all AF2 models are apo structures. On the other hand, the worst match is given by FLT3, predicted in the CIDO state (Fig. 3C). CIDI structures are the most represented conformational state present in the PDB (Fig. 1), while CIDO are the second last in terms of representation. Reviewing the models, we observe that AF2 reproduces the state-specific arrangement of binding site residues needed to recognize active molecules for the CIDI conformation, while the poorest models can be observed for the CIDO conformation (Fig. 3D-F).

**Figure 3:**
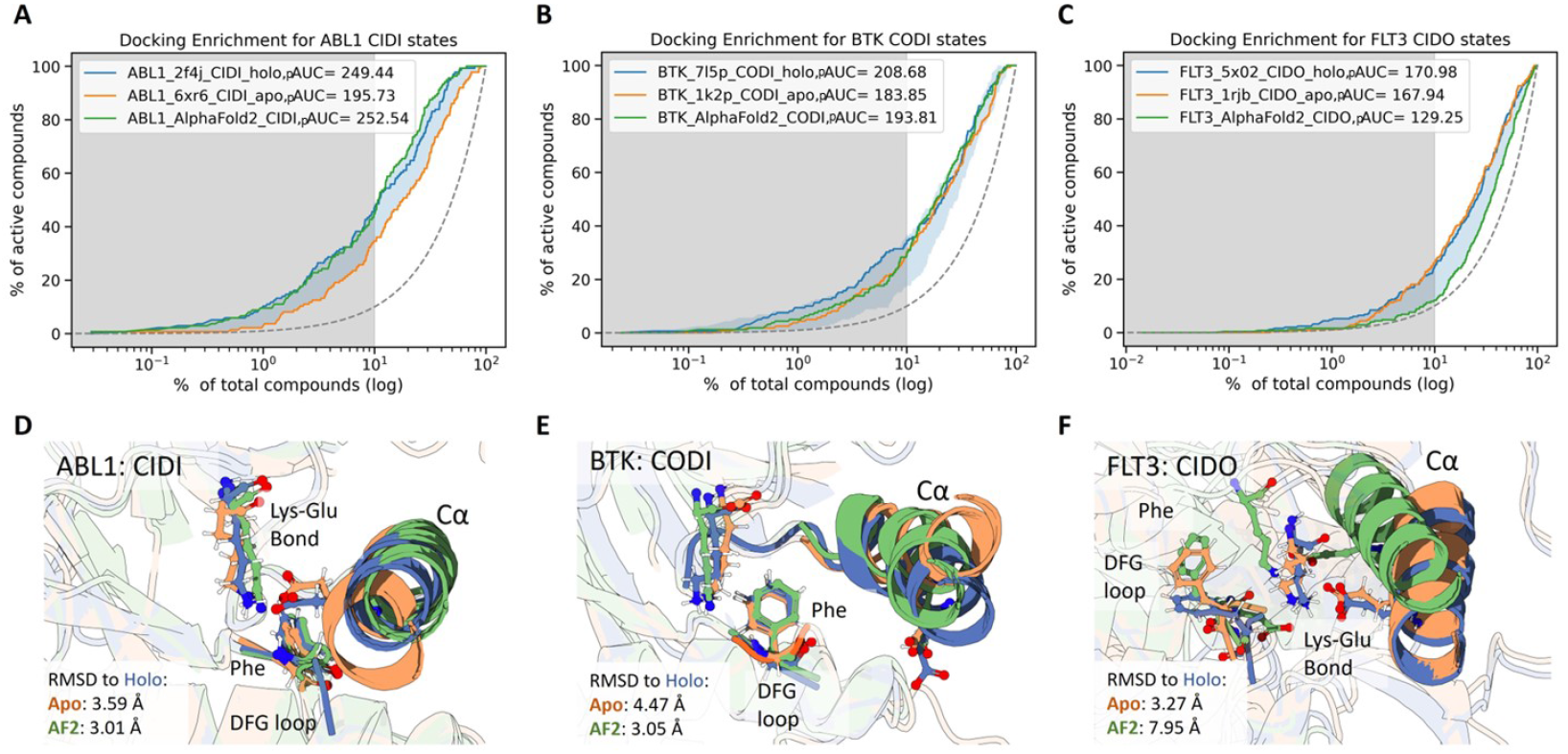
Docking enrichment curves for AF2-generated structures versus minimized crystal representatives. (A-C) Docking enrichment curves for ABL1 (CIDI), BTK (CODI), FLT3 (CIDO). Blue lines represent enrichments for highest scoring structure, orange lines represent enrichments for apo structures, and green lines describe AF2 models. The blue shaded region represents the range of enrichment scores observed for all crystal structures. (D-F) Structural superimposition of the c*α*-Helix, DFG, and Lys-Glu hydrogen bond for ABL1 (CIDI), BTK (CODI), FLT3 (CIDO). The RMSD values were measured for theses structural elements with respect to the holo counterparts.

Finally, we evaluated the performance of AF2 models in maintaining known kinase-ligand interactions in the binding pocket (Fig. 4). Compared to both redocking and cross docking experiments of crystal ligands to conformation-matched structures of kinases, AF2 models consistently perform the poorest in recapitulating all known kinase-ligand interactions.

**Figure 4:**
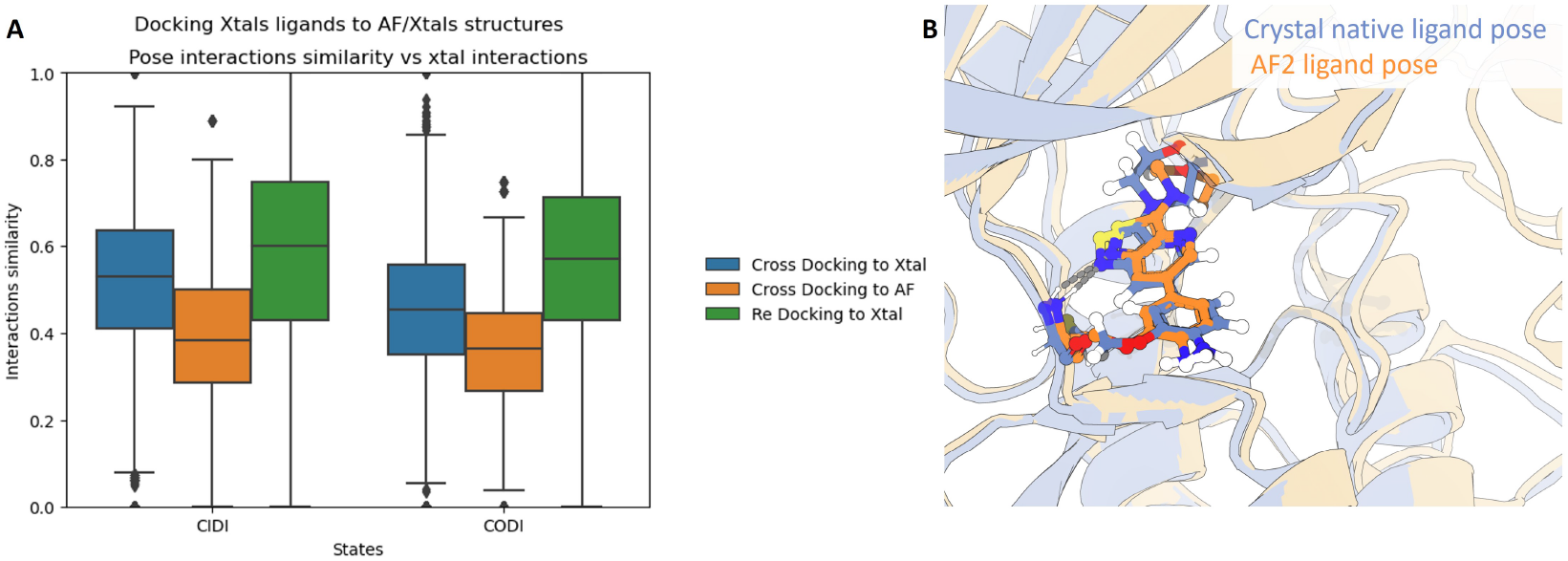
A) Evaluation of kinase-ligand binding site interactions of AF2 structures. Ligands co-crystallized with kinases in the CIDI and CODI conformations where re-docked into their original structure, or cross docked into a conformation-matched crystal structure or the respective AF2 model. Interaction similarity is defined as the Jaccard coefficient between the interactions observed in the docked structures to the interactions observed in the native structure. B) Pose of highest performing AF2 model in terms of interaction accuracy vs native ligand crystal pose.

Taken together, in case of high representation of a conformational state in the PDB, the AF2 structures provide comparable early enrichment in the identification of true actives in the docking benchmark set compared to the minimized crystal structures of the same conformational state (Fig. 3). On the other hand, AF2 structures may lose key interactions that facilitate ligand binding which may directly impact the prioritization molecules during activities such as virtual screening. Critically, the amount of representation of a given conformational state in the PDB correlates with enrichment performance, and thus the quality of the generated structure.

## 4 Discussion

Elucidating the conformational landscape of protein kinases may lead to key insights into cellular signalling mechanisms as well as enable the discovery of more effective therapeutics. We investigated the capability of AF2 to model protein kinases across several conformational states. Given that the majority of kinase structures in the PDB are in the active conformation (Fig. 1A), we initially hypothesized that AF2 may confidently model just the active conformation of kinases. Surprisingly, AF2 is able to model several conformational states of protein kinases, and certain conformations are observed in specific kinase families (Fig. 1B). This observation is due to CIDO, CODI, CODO, and *ω*CD conformations often being stabilized with a ligand bound (7). Thus, kinase families enriched with known drug targets are privileged in terms of conformational diversity in both native crystal and AF2-modeled structures. Importantly, the AF2 models maintained key evolutionary and structural relationships between kinases (Fig. 1C).

We then investigated the confidence of AF2 predictions in the 3D positioning of motifs that determine kinase conformations. We observed that at these motifs pLDDT reveals significant variability in the confidence of these coordinates (Fig. 2). The high variability of pLDDT at these residues is seen for kinases that have multiple solved crystal structures in several conformations. We propose that the variability of pLDDT in AF2 models may, in part, be explained by the conformational diversity of individual kinases appearing during the training process.

Finally, we performed docking against both AF2 and conformation-matched protein kinase crystal structures to evaluate the utility of AF2 models in virtual screening. In early enrichment of known ligands, the AF2 models performed comparably to native crystal structures across conformational states.

Recently, several studies have evaluated the utility of AF2 models for structure based drug discovery (SBDD). Recent studies by Zhang et al.(37) and Rovira et al.(38) have shown that certain AF2 models need structural refinement before being used in virtual screening campaigns. Whilst this observation might hold from a global view of the entire protein landscape, we observe that integrating protein conformational dynamics is critical for proper evaluation of AF2 models in SBDD. We observe that differences arise when analyzing the conformational states of AF2 kinase structures separately (Fig. 3). Specifically, that the AF2 model of ABL1 kinase in the active state performed comparably to the best holo crystal structure. Furthermore, we observed that the binding site residues of this predicted structure were accurately modeled by AF2 to match a holo active structure. On the other hand, loss in performance was only seen in the CIDO structure of FLT3. We posit that this is due to the lack of CIDO representation during the AF2 training process, thus having an impact on the quality of the generated structure.

We conclude that AF2-modeled kinase structures can be used to effectively model the conformational landscape of the kinase active site for highly represented conformational states. We see a large opportunity to leverage AF2 to rationally design novel, conformation-specific inhibitors for kinases lacking solved structures.

## References

[1] A. A. Zarrin, K. Bao, P. Lupardus, and D. Vucic, “Kinase inhibition in autoimmunity and inflammation,” Nature Reviews Drug Discovery, vol. 20, no. 1, pp. 39–63, 2021.

[2] F. M. Ferguson and N. S. Gray, “Kinase inhibitors: the road ahead,” Nature reviews Drug discovery, vol. 17, no. 5, pp. 353–377, 2018.

[3] P. Cohen, “Protein kinases—the major drug targets of the twenty-first century?,” Nature reviews Drug discovery, vol. 1, no. 4, pp. 309–315, 2002.

[4] R. Röck, J. E. Mayrhofer, O. Torres-Quesada, F. Enzler, A. Raffeiner, P. Raffeiner, A. Feichtner, R. G. Huber, S. Koide, S. S. Taylor, J. Troppmair, and E. Stefan, “BRAF inhibitors promote intermediate BRAF(v600e) conformations and binary interactions with activated RAS,” vol. 5, no. 8, p. eaav8463. Publisher: American Association for the Advancement of Science.

[5] A. Haldane, W. F. Flynn, P. He, R. S. K. Vijayan, and R. M. Levy, “Structural propensities of kinase family proteins from a Potts model of residue co-variation,” Protein Science: A Publication of the Protein Society, vol. 25, no. 8, pp. 1378–1384, 2016.

[6] P. M.-U. Ung, R. Rahman, and A. Schlessinger, “Redefining the Protein Kinase Conformational Space with Machine Learning,” Cell Chemical Biology, vol. 25, pp. 916–924.e2, July 2018.

[7] R. Roskoski Jr, “Classification of small molecule protein kinase inhibitors based upon the structures of their drug-enzyme complexes,” Pharmacological research, vol. 103, pp. 26–48, 2016.

[8] J. L. Sussman, D. Lin, J. Jiang, N. O. Manning, J. Prilusky, O. Ritter, and E. E. Abola, “Protein data bank (pdb): database of three-dimensional structural information of biological macromolecules,” Acta Crystallographica Section D: Biological Crystallography, vol. 54, no. 6, pp. 1078–1084, 1998.

[9] V. Modi and R. L. Dunbrack, “Defining a new nomenclature for the structures of active and inactive kinases,” Proceedings of the National Academy of Sciences, vol. 116, pp. 6818–6827, Apr. 2019. Company: National Academy of Sciences Distributor: National Academy of Sciences Institution: National Academy of Sciences Label: National Academy of Sciences Publisher: Proceedings of the National Academy of Sciences.

[10] J. Jumper, R. Evans, A. Pritzel, T. Green, M. Figurnov, O. Ronneberger, K. Tunyasuvunakool, R. Bates, A. Žídek, A. Potapenko, A. Bridgland, C. Meyer, S. A. A. Kohl, A. J. Ballard, A. Cowie, B. Romera-Paredes, S. Nikolov, R. Jain, J. Adler, T. Back, S. Petersen, D. Reiman, E. Clancy, M. Zielinski, M. Steinegger, M. Pacholska, T. Berghammer, S. Bodenstein, D. Silver, O. Vinyals, A. W. Senior, K. Kavukcuoglu, P. Kohli, and D. Hassabis, “Highly accurate protein structure prediction with AlphaFold,” vol. 596, no. 7873, pp. 583–589. Number: 7873 Publisher: Nature Publishing Group.

[11] D. del Alamo, D. Sala, H. S. Mchaourab, and J. Meiler, “Sampling alternative conformational states of transporters and receptors with AlphaFold2,” eLife, vol. 11, p. e75751, Mar. 2022. Publisher: eLife Sciences Publications, Ltd.

[12] M. Varadi, S. Anyango, M. Deshpande, S. Nair, C. Natassia, G. Yordanova, D. Yuan, O. Stroe, G. Wood, A. Laydon, et al., “Alphafold protein structure database: massively expanding the structural coverage of protein-sequence space with high-accuracy models,” Nucleic acids research, vol. 50, no. D1, pp. D439–D444, 2022.

[13] P. D. Bank, “Protein data bank,” Nature New Biol, vol. 233, p. 223, 1971.

[14] J.-M. Chandonia, N. K. Fox, and S. E. Brenner, “Scope: classification of large macromolecular structures in the structural classification of proteins—extended database,” Nucleic acids research, vol. 47, no. D1, pp. D475–D481, 2019.

[15] U. Consortium, “Uniprot: a hub for protein information,” Nucleic acids research, vol. 43, no. D1, pp. D204–D212, 2015.

[16] R. Rahman, P. M.-U. Ung, and A. Schlessinger, “KinaMetrix: a web resource to investigate kinase conformations and inhibitor space,” Nucleic acids research, vol. 47, no. D1, pp. D361–D366, 2019. Publisher: Oxford University Press.

[17] V. Modi and R. L. Dunbrack, “A structurally-validated multiple sequence alignment of 497 human protein kinase domains,” vol. 9, no. 1, p. 19790.

[18] R. C. Edgar, “Muscle: multiple sequence alignment with high accuracy and high throughput,” Nucleic acids research, vol. 32, no. 5, pp. 1792–1797, 2004.

[19] L. Schrödinger and W. DeLano, “Pymol.”

[20] A. J. Kooistra, G. K. Kanev, O. P. van Linden, R. Leurs, I. J. de Esch, and C. de Graaf, “Klifs: a structural kinase-ligand interaction database,” Nucleic acids research, vol. 44, no. D1, pp. D365–D371, 2016.

[21] P. J. Cock, T. Antao, J. T. Chang, B. A. Chapman, C. J. Cox, A. Dalke, I. Friedberg, T. Hamelryck, F. Kauff, B. Wilczynski, et al., “Biopython: freely available python tools for computational molecular biology and bioinformatics,” Bioinformatics, vol. 25, no. 11, pp. 1422–1423, 2009.

[22] D. Sydow, E. Aßmann, A. J. Kooistra, F. Rippmann, and A. Volkamer, “Kissim: Predicting off-targets from structural similarities in the kinome,” Journal of Chemical Information and Modeling, vol. 62, no. 10, pp. 2600–2616, 2022.

[23] L. Van der Maaten and G. Hinton, “Visualizing data using t-sne.,” Journal of machine learning research, vol. 9, no. 11, 2008.

[24] V. Le Guilloux, P. Schmidtke, and P. Tuffery, “Fpocket: an open source platform for ligand pocket detection,” BMC bioinformatics, vol. 10, no. 1, pp. 1–11, 2009.

[25] D. Sydow, J. Rodríguez-Guerra, and A. Volkamer, “Opencadd-klifs: A python package to fetch kinase data from the klifs database,” Journal of Open Source Software, vol. 7, no. 70, p. 3951, 2022.

[26] M. Ester, H.-P. Kriegel, J. Sander, and X. Xu, “A Density-based Algorithm for Discovering Clusters a Density-based Algorithm for Discovering Clusters in Large Spatial Databases with Noise,” in Proceedings of the Second International Conference on Knowledge Discovery and Data Mining, KDD’96, (Portland, Oregon), pp. 226–231, AAAI Press, 1996.

[27] C. C. Group, “Molecular operating environment,” 2018.

[28] G. Landrum et al., “Rdkit: A software suite for cheminformatics, computational chemistry, and predictive modeling,” Greg Landrum, 2013.

[29] F. Imrie, A. R. Bradley, and C. M. Deane, “Generating property-matched decoy molecules using deep learning,” Bioinformatics, vol. 37, no. 15, pp. 2134–2141, 2021.

[30] N. M. O’Boyle, M. Banck, C. A. James, C. Morley, T. Vandermeersch, and G. R. Hutchison, “Open babel: An open chemical toolbox,” Journal of cheminformatics, vol. 3, no. 1, pp. 1–14, 2011.

[31] D. R. Koes, M. P. Baumgartner, and C. J. Camacho, “Lessons learned in empirical scoring with smina from the csar 2011 benchmarking exercise,” Journal of chemical information and modeling, vol. 53, no. 8, pp. 1893–1904, 2013.

[32] S. Salentin, S. Schreiber, V. J. Haupt, M. F. Adasme, and M. Schroeder, “Plip: fully automated protein–ligand interaction profiler,” Nucleic acids research, vol. 43, no. W1, pp. W443–W447, 2015.

[33] V. Modi and R. L. Dunbrack, Jr, “Kincore: a web resource for structural classification of protein kinases and their inhibitors,” vol. 50, pp. D654–D664.

[34] J. L. Binder, J. Berendzen, A. O. Stevens, Y. He, J. Wang, N. V. Dokholyan, and T. I. Oprea, “AlphaFold illuminates half of the dark human proteins,” vol. 74, p. 102372.

[35] T. Hegedűs, M. Geisler, G. L. Lukács, and B. Farkas, “Ins and outs of alphafold2 transmembrane protein structure predictions,” Cellular and Molecular Life Sciences, vol. 79, no. 1, pp. 1–12, 2022.

[36] H.-B. Guo, A. Perminov, S. Bekele, G. Kedziora, S. Farajollahi, V. Varaljay, K. Hinkle, V. Molinero, K. Meister, C. Hung, et al., “Alphafold2 models indicate that protein sequence determines both structure and dynamics,” 2022.

[37] P. Zhang, Z. Wei, C. Che, and B. Jin, “DeepMGT-DTI: Transformer network incorporating multilayer graph information for Drug–Target interaction prediction,” Computers in Biology and Medicine, p. 105214, Jan. 2022.

[38] A. M. Diaz-Rovira, H. Martin, T. Beuming, L. Diaz, V. Guallar, and S. S. Ray, “Are deep learning structural models sufficiently accurate for virtual screening? application of docking algorithms to alphafold2 predicted structures,” bioRxiv, 2022.

